# Geometry-Controlled Diffusion in Bicontinuous Cubic Lipid Networks: Network Topology, Tortuosity, and Pore-Scale Constriction Effects

**DOI:** 10.64898/2026.01.14.699444

**Authors:** Amul Ojha, Arya Ojha, Adarsh Sharma, Samiksha Mesharam

## Abstract

Triply periodic minimal surface (TPMS) architectures—Gyroid, Diamond, and Primitive—are increasingly used as delivery matrices for therapeutic agents. Although point-tracer diffusion and geodesic tortuosity (*τ*) describe solvent transport through their aqueous nanochannels, practical drug delivery requires accounting for finite-sized solutes. We present a unified computational framework that combines discrete lattice random walks with periodic Euclidean distance transforms and domain erosion. By mapping local clearance from lipid boundaries, we quantify both the effective diffusion coefficient (*D*_eff_) and the macroscopic size-dependent sieving factor *K*_*S*_ (*ρ*) over a range of normalized tracer radii (*ρ* = *r*/ *R*_ch_). Our results decouple structural tortuosity from excluded-volume partition effects, providing a rigorous 3D generalization of classical hindrance theory tailored to the complex topology of TPMS matrices. Comparison with experimental water self-diffusion data shows that the geometric point-tracer model systematically overpredicts the observed diffusivity; the residual offset is consistent with a bound-water contribution, though we do not claim a quantitative scaling law (the correlation rests on only five data points and is reported descriptively).All simulations are reported in dimensionless units; a physical length scale is required to map normalized sieving to absolute drug radii. Using a typical monoolein gyroid lattice constant (*a* ≈ 10 nm) as an illustrative conversion, water’s hydrodynamic radius corresponds to *ρ* ≈ 0.12 (with the Marching Cubes-corrected *R*_ch_), where steric hindrance is negligible (*K*_*S*_ ≈ 1), whereas drug-sized payloads (*ρ* ≳ 0.3) experience substantial steric exclusion.

## Introduction

Amphiphilic molecules spontaneously assemble into bicontinuous cubic lipid phases that separate a continuous bilayer into two interpenetrating aqueous networks with the symmetry of the gyroid 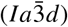, diamond 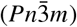, or primitive 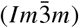 triply periodic minimal surfaces (TPMS). ^1,15,29^ Dispersions of these phases—cubosomes—offer a large internal surface area, mechanical stability, and the ability to co-encapsulate both hydrophilic and hydrophobic cargos, making them promising candidates for controlled drug delivery. ^3,11,23,30^

Molecular transport through these networks is characterized by effective diffusion coefficients *D*_eff_ that are significantly lower than the bulk value *D*_0_, as measured by pulsed-field gradient NMR (PFG-NMR) and electrochemistry. Anderson and Wennerström ^2^ developed obstruction-factor theory for diffusion on diamond and Schwarz-P IPMS structures and compared it with early experimental data. Eriksson and Lindblom ^7^ later reported PFG-NMR self-diffusion measurements for monoolein/water and monolinolein/water systems across the gyroid and diamond cubic phase regions. More recent electrochemical and PFG-NMR studies using larger, drug-sized tracers ^19,22^ reveal an additional, size-dependent steric contribution that standard point-tracer obstruction theories cannot capture.

Idealized point-tracer models attribute the reduction in *D*_eff_ solely to the tortuosity *τ* of the aqueous pathways. ^32^ While this approach works for small tracers such as water molecules, it fails to predict the behavior of larger, finite-sized solutes.

The three TPMS geometries have distinct connectivities: gyroid features smoothly curved, three-coordinated junctions; diamond has tetrahedral, four-coordinated junctions; and primitive has six-coordinated junctions, though it is more prone to network disconnection at low hydration despite having larger pore bodies. ^1,9,10^(We note that “Schwarz-P IPMS” is the legacy name for the primitive minimal surface, which is the same family of triply periodic structures studied here.)

To address these issues, we: (*i*) simulate point-tracer diffusion directly on all three TPMS level-set geometries and extract *D*_eff_ / *D*_0_ from the long-time mean-squared displacement (MSD); (*ii*) compute geodesic tortuosity on the same discretized network to ensure consistency between *D*_eff_ and *τ*; (*iii*) compare the resulting point-tracer predictions with open-access experimental water self-diffusion data, quantifying the residual gap; and (*iv*) introduce a size-dependent sieving factor *K*_*S*_ (*ρ*) using periodic Euclidean distance transforms and domain erosion—a theoretical construct that awaits experimental validation for drug-sized solutes.

## Model and Methods

### TPMS geometry and lattice random-walk simulation

We approximate each TPMS by the standard trigonometric (Fourier) level-set field *f* (***r***) : *f*_*G*_ = sin *x* cos *y*+ sin *y* cos *z*+ sin *z* cos *x, f*_*D*_ = sin *x* sin *y* sin *z*+ sin *x* cos *y* cos *z* +cos *x* sin *y* cos *z*+ cos *x* cos *y* sin *z, f*_*P*_ = cos *x*+ cos *y* +cos *z*, with *x, y, z* ∈[0, 2π) spanning one cubic unit cell of edge *a*. ^1,10,14^ The aqueous domain is {*f* ***r*** > *t*}; because these fields are approximately inversion-symmetric, the total water volume fraction is *ϕ*_*w*_ ≈2 vol (*f* > *t*), and *t* (*ϕ*_*w*_) is obtained by numerical inversion on a discrete grid. ^21^ The only inputs are the trigonometric fields, the target *ϕ*_*w*_, and the lattice size *N* = 100; the threshold *t* is uniquely determined, with no calibration parameters.

We model diffusion as a lattice random walk that mimics Brownian dynamics with reflecting boundaries. On an *N*^3^ periodic grid (*N* = 100), tracers attempt unit steps to one of six face-neighbouring voxels chosen uniformly at random. ^32^ Steps that enter a lipid voxel are rejected, leaving the tracer stationary for that time step; unwrapped displacements are accumulated for the MSD calculation. ^8,20^ The effective diffusion coefficient *D*_eff_ is extracted from the long-time linear slope of the ensemble-averaged MSD. For point-tracer and hydration-sweep simulations, we use *N*_walkers_ = 5000, *N*_steps_ = 6000, and 10 independent seeds. For size-dependent sieving, the larger parameter space leads us to reduce the ensemble to *N*_walkers_ = 2000, *N*_steps_ = 3000, and 5 seeds. A convergence check at one common condition (gyroid, *ϕ*_*w*_ = 0.35, *ρ* = 0) shows that the reduced ensemble differs from the full one by less than 2% in *D*_eff_ / *D*_0_, confirming the adequacy of the smaller ensemble. The open-grid reference *D*_0_ is computed under the same settings; the ratio *D*_eff_ / *D*_0_ is therefore independent of absolute length or time scales.

### Lattice connectivity and discretisation sensitivity

We assess the influence of grid anisotropy by comparing the standard 6-connected nearest-neighbour scheme with 18- and 26-connected moves (which include face, edge, and corner-neighbouring voxels). For the gyroid phase at *ϕ*_*w*_ = 0.35, the 6-connected scheme gives *D*_eff_ /*D*_0_ = 0.461 ± 0.008, while the 18- and 26-connected schemes yield 0.478 ± 0.010 and 0.483 ± 0.011, respectively. The corresponding tortuosity values are *τ* = 1.544 ± 0.126 (6-connected), 1.511± 0.118 (18-connected), and 1.498± 0.115 (26-connected). These differences are modest (< 5% for *D*_eff_ /*D*_0_ and < 3% for *τ*) and do not alter the reported trends or the relative ordering of the phases. We choose the 6-connected scheme for consistency with most voxel-based diffusion studies and to avoid biasing the distance transforms, which are also 6-connected.

#### MSD fitting and diffusive regime analysis

We extract the MSD from the linear regime of the ensemble-averaged mean-squared displacement versus time. For point-tracer and hydration-sweep simulations, the fitting window covers the final 50% of the trajectory (*N*_steps_ = 3000 to 6000), after verifying that the MSD has crossed from the initial ballistic/transient regime into the asymptotic diffusive regime. The crossover time is defined by MSD (*t*) > 4 (in voxel^2^), corresponding to *t* ≳ 1000 steps for all phases and hydrations. The coefficient of determination *R*^2^ for the linear fits exceeds 0.999 in all cases. For the size-dependent sieving simulations (with reduced *N*_steps_ = 3000), we apply the same criterion and use the fitting window from *N*_steps_ / 2 to *N*_steps_ (*R*^2^ > 0.998). We observe no subdiffusive transients near percolation or for large *ρ*; when the eroded domain approaches the percolation threshold, we compute the MSD slope only for percolating configurations and report the percolation status separately (Figure 4).

We identify connected components of the aqueous domain using a periodic 6-connected cluster-labeling algorithm. Refining the grid to *N* = 150 shifts the estimated percolation threshold by less than 0.02 in *ϕ*_*w*_, confirming that the disconnection is not a lattice artifact. We do not attempt to pinpoint the precise primitive-phase percolation threshold because that would require a dedicated finite-size scaling study; instead, we benchmark all primitive-phase results at *ϕ*_*w*_ = 0.45, a hydration that our cluster-labeling analysis confirms to yield a fully connected single labyrinth well above the disconnection boundary.

The Euclidean distance transform and geodesic distances are evaluated on the same 6-connected periodic lattice, with the latter obtained using Dijkstra’s shortest-path algorithm.

For qualitative visualization, we export selected configurations and render them using OVITO (Open Visualization Tool). ^31^ In these renderings, the lipid matrix appears in off-white and tracer particles in black.

### Geodesic tortuosity

We compute tortuosity on the *same* voxel connectivity graph used for the random-walk simulation (6-connected, periodic) using multi-source Dijkstra shortest-path algorithms from 20–25 randomly selected source voxels to all target destinations. For each source, *τ* is the ratio of the geodesic path length to the Euclidean distance for target voxels at intermediate separations (0.30–0.48 of the box length). ^13,32^ This window avoids both trivial short-path bias and long-range periodic-image artifacts. ^16^ Using the same graph for both *D*_eff_ and *τ* ensures full consistency between the two transport descriptors.

### Size-dependent sieving

To account for the finite hydrodynamic size of therapeutic payloads, we determine the accessible aqueous region for a spherical tracer of radius *r* by eroding the TPMS pore network. Specifically, we exclude water voxels whose Euclidean distance to the nearest lipid boundary—computed via a periodic Euclidean distance transform—is less than or equal to *r*. We then run the same random-walk protocol on each eroded domain to obtain *D*_eff_ (*r*).

The reference channel radius *R*_ch_ is defined operationally as twice the hydraulic radius of the uneroded aqueous labyrinth:

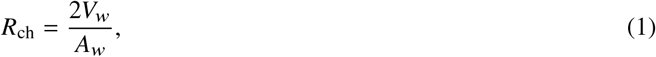

where *V*_*w*_ is the volume of the aqueous labyrinth and *A*_*w*_ is its interfacial area with the lipid bilayer. To reduce the known resolution sensitivity of voxel-based area estimators, we evaluate *A*_*w*_ using a Marching Cubes surface reconstruction of the level-set field *f* (***r***) at the threshold *t* for the target *ϕ*_*w*_. ^18^ This approach reduces the systematic bias in *A*_*w*_ (and hence *R*_ch_) to < 5% for *N* ≥ 100, as confirmed by resolution convergence tests. The Marching Cubes-derived *R*_ch_ values are 15.19 voxels for gyroid, 12.50 for diamond, and 23.87 for primitive. All reported *ρ* = *r* /*R*_ch_ values and *K*_*S*_ (*ρ*) curves use these corrected values. This definition is topology- and hydration-specific; *R*_ch_ is not universal across phases.

Currently, *R*_ch_ is expressed in voxel units. Converting *ρ* to an absolute tracer radius in nanometers requires the physical lattice constant *a* of the specific lipid/water system. We leave *a* unspecified because our simulations are intended to be general across cubic phase compositions; consequently, the normalized sieving results are dimensionless. We discuss this limitation further in the Limitations section.

For illustration, we can use a typical monoolein-based gyroid lattice constant *a* ≈ 10 nm (consistent with reported values for hydrated monoolein cubic phases, typically ~ 10–14 nm). ^26^ At *N* = 100, one voxel edge is *a* /*N* ≈ 0.1 nm, so *R*_ch_ = 15.19 voxels corresponds to about 1.52 nm, and *ρ* = 0.20 corresponds to *r* ≈ 0.30 nm. This is only an illustrative example, not a validated calibration; different lipid systems and hydrations will have different lattice constants. The uncorrected voxel-based *R*_ch_ = 9.24 voxels would have given *R*_ch_ ≈ 0.92 nm, so the Marching Cubes correction systematically increases the physical length scale by a factor of ~ 1.6.

The accessible volume fraction is

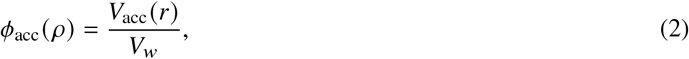

where *V*_acc_ (*r*) is the number of aqueous voxels remaining after erosion and *V*_*w*_ is the total number of aqueous voxels before erosion.

The macroscopic sieving factor *K*_*S*_ (*ρ*) is the product of the local mobility restriction and the steric partitioning coefficient:

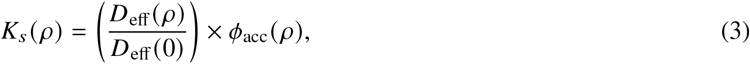

where *D*_eff_ (0) is the point-tracer diffusivity in the uneroded network. This product form follows from a simple homogenisation argument: for a periodic porous medium with accessible volume fraction *ϕ*_acc_, the macroscopic effective diffusivity is the product of the volume fraction and the diffusivity within the accessible pore space, provided that the accessible pores are well-connected and partition equilibrium holds at the macroscale. ^32^ Here, *D*_eff_ (*ρ*) /*D*_eff_ (0) accounts for excluded-volume reduction in mobility, while *ϕ*_acc_ (*ρ*) captures the reduction in transport volume. This construction separates local mobility restriction from steric partitioning and provides a 3D generalization of classical steric hindrance and entropy-barrier theories ^6,28,33^ adapted to the non-idealized topology of TPMS architectures.

### Experimental data

To test the predictive power of the geometric point-tracer model, we compare our simulations with the open-access PFG-NMR data of Eriksson and Lindblom ^7^ for the monoolein/water system at 25 °C (Table 1). This dataset gives experimental water self-diffusion coefficients 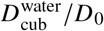 along with the fraction of bound water *p*_B_ from a two-site exchange model.

**Table 1.** Comparison of experimental PFG-NMR data and geometric point-tracer simulations for the monoolein/water system at 25 °C (experimental data adapted from Eriksson and Lindblom ^7^). Aqueous volume fractions *ϕ*_*w*_ are calculated using *ρ*_lipid_ = 1.075 g cm^−3^ (*v*_lipid_ = 0.93 cm^3^ g^−1^) ^27^. Simulation values are mean ± standard deviation over ten independent random-walk seeds. Here *τ*_app_ = *ϕ*_*w*_ / (*D*_cub_ /*D*_0_) _exp_ is the apparent tortuosity derived from experiment, distinct from the geodesic tortuosity *τ* calculated from simulation. Also shown for comparison are experimental diffusion data for doxorubicin and K_4_Fe(CN)_6_ from Nazaruk et al. ^22^ (bottom rows).

| Phase Type | Water wt% | $\phi_w$ | $p_B$ (Exp)<sup>7</sup> | $(D_{\text{cub}}/D_0)_{\text{exp}}$<sup>7</sup> | $\tau_{\text{app}}$ | $(D_{\text{eff}}/D_0)_{\text{sim}}$ |
| --- | --- | --- | --- | --- | --- | --- |
| Gyroid ($Ia\bar{3}d$) | 28.0 | 0.295 | 0.56 | 0.201 | 1.47 | $0.443 \pm 0.013$ |
| Gyroid ($Ia\bar{3}d$) | 30.4 | 0.320 | 0.49 | 0.237 | 1.35 | $0.455 \pm 0.009$ |
| Gyroid/Diamond ($G/D$) | 33.4 | 0.350 | 0.41 | 0.276 | 1.27 | $0.461 \pm 0.008$ (G)<br>$0.413 \pm 0.012$ (D) |
| Diamond ($Pn\bar{3}m$) | 36.6 | 0.383 | 0.39 | 0.292 | 1.31 | $0.425 \pm 0.011$ |
| Diamond ($Pn\bar{3}m$) | 39.5 | 0.412 | 0.38 | 0.300 | 1.37 | $0.442 \pm 0.015$ |
| *Electrochemical diffusion data (Nazaruk et al.<sup>22</sup>)* |  |  |  |  |  |  |
| Solute | Phase | Buffer | $r$ (nm) | $\rho$ | $D_{\text{eff}}/D_0$ (exp) | $K_s$ (predicted) |
| Doxorubicin | MO gyroid | MES (pH 5.5) | 0.6 | 0.39 | 0.48 | 0.27 |
| Doxorubicin | MO gyroid | HEPES (pH 7.5) | 0.6 | 0.39 | 0.51 | 0.27 |
| $\text{K}_4\text{Fe}(\text{CN})_6$ | MO gyroid | $\text{H}_2\text{O}$ (0.1M KCl) | 0.3 | 0.20 | 0.20–0.26 | 0.56 |

We compute aqueous volume fractions *ϕ*_*w*_ using *ρ*_lipid_ = 1.075 g cm^−3^ (*v*_lipid_ = 0.93 cm^3^ g^−1^), ^27^ matching the original study. Experimental apparent tortuosities are *τ*_app_ = *ϕ*_*w*_ / (*D*_eff_ / *D*_0_) _exp_. The lattice/time normalization cancels in *D*_eff_ / *D*_0_; the threshold-to-*ϕ*_*w*_ mapping is fixed solely by the level-set volume fraction, so no external calibration or adjustable parameters enter the comparison. ^12^ We use all five 25 °C data points covering the gyroid and diamond phases.

## Results and Discussion

**Figure 1.**
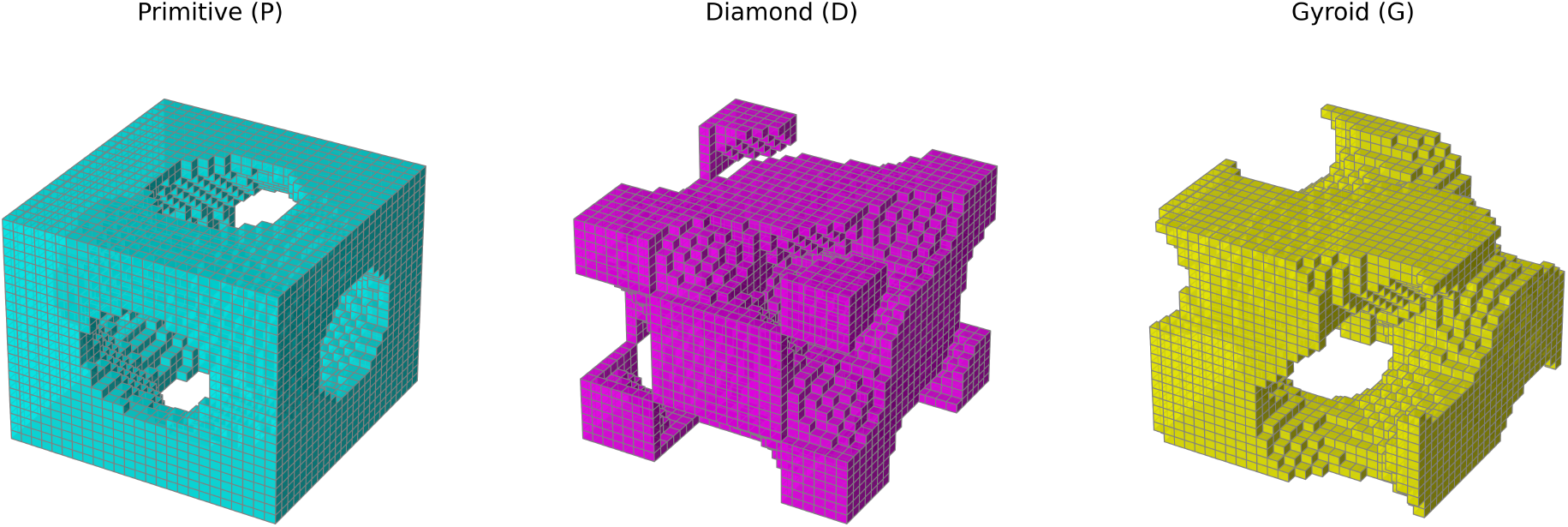
Trigonometric level-set approximations of TPMS cubic phase architectures. High-resolution 3D voxel representations at *N* = 25 (illustrative only; simulations use *N* = 100) of the single labyrinth aqueous domains at the physical water volume fraction *ϕ*_*w*_ = 0.35 (threshold *t* corresponding to finite bilayer thickness) for the (left) Primitive 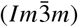, (middle) Diamond 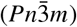, and (right) Gyroid 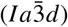 structures. The distinct channel networks and junction connectivities govern both the geodesic tortuosity (*τ*) and the size-dependent steric sieving behavior of encapsulated solutes.

### Transport hierarchy and tortuosity

Our connectivity analysis shows that gyroid and diamond single labyrinths remain continuously percolating throughout the studied hydration range, whereas the primitive single labyrinth fragments at low hydration. Because the primitive single labyrinth loses connectivity below its percolation threshold (*ϕ*_*w*_ ≈ 0.40), it cannot be evaluated at *ϕ*_*w*_ = 0.35. To compare all three architectures in their fully percolating, open-channel regimes, we evaluate gyroid and diamond at *ϕ*_*w*_ = 0.35 and primitive at its lowest connected hydration, *ϕ*_*w*_ = 0.45. Once percolating, the primitive phase’s transport is dominated by constricted inter-pore necks rather than pore-body dimensions.

From the long-time MSD slopes, we obtain 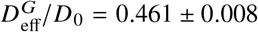 and 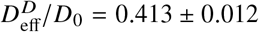 at *ϕ*_*w*_ = 0.35, and 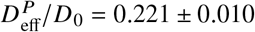 at *ϕ*_*w*_ = 0.45. At equal hydration, gyroid diffuses faster than diamond (0.461 vs. 0.413). Remarkably, primitive remains substantially more hindered than both gyroid and diamond at *ϕ*_*w*_ = 0.35, despite being granted a higher water content (*ϕ*_*w*_ = 0.45). Thus, we do not claim a universal fixed-hydration hierarchy; the robust finding is that primitive is the most restrictive once connected, and among gyroid and diamond at equal hydration, gyroid is faster.

The corresponding geodesic tortuosity distributions give 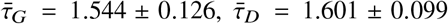 and 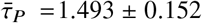. The tortuosity intervals overlap substantially, so the primitive phase’s severe transport deficit is not explained by path length; rather, it reflects the dominance of bottleneck constrictions.

The OVITO snapshots in Figure 2 qualitatively illustrate the different channel geometries and the spatial distribution of tracer particles. In gyroid, particles populate smooth, three-coordinated channels; in diamond, they reside at tetrahedrally connected nodes; and in primitive, they occupy a more open but bottleneck-limited network. These visual differences are consistent with the quantitative diffusion coefficients.

**Figure 2.**
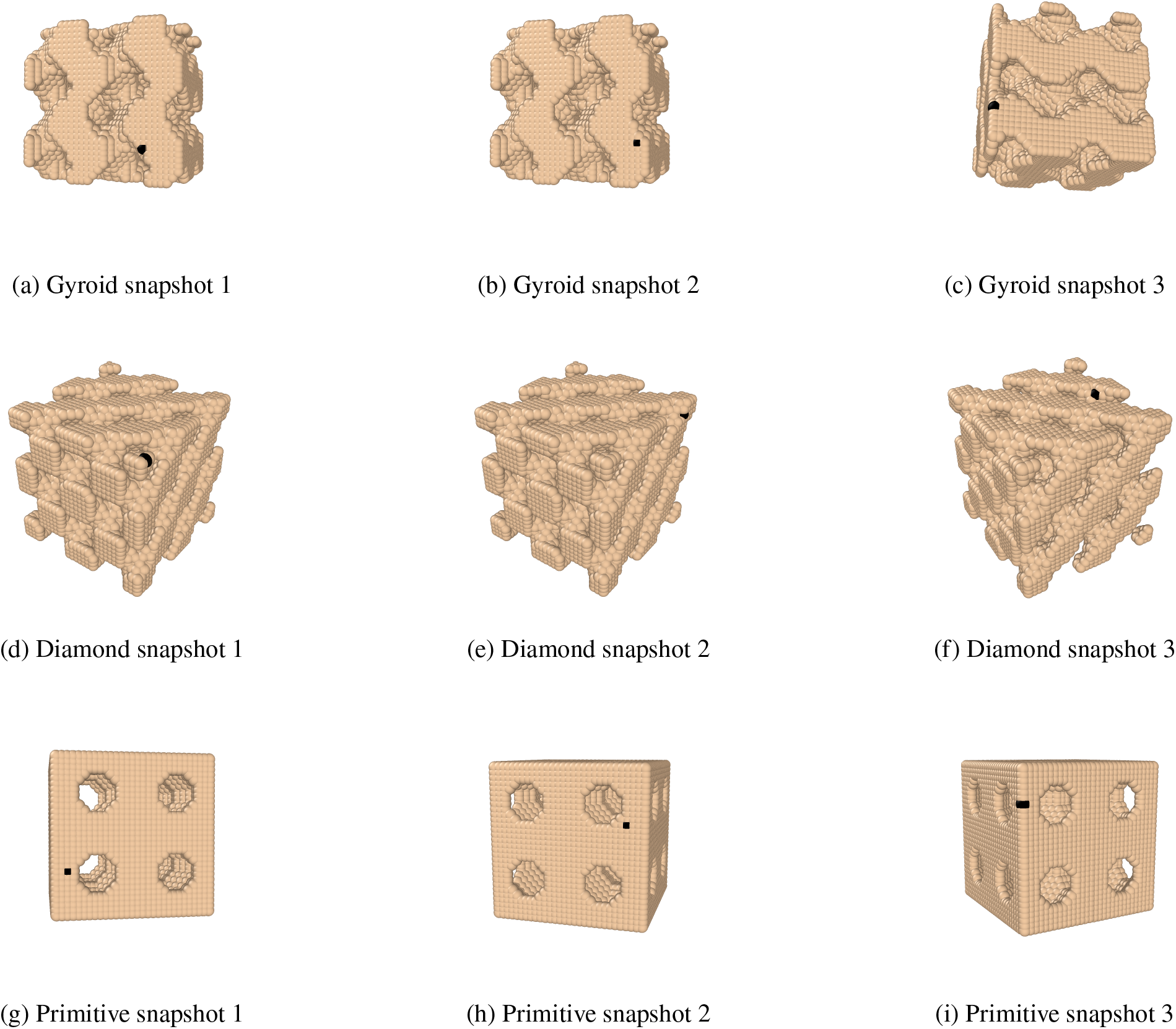
Representative OVITO renderings of instantaneous tracer configurations in the three TPMS phases. Top row: gyroid 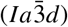; middle row: diamond 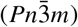; bottom row: primitive 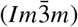. Lipid matrix is shown in off-white; tracer particles are black. The panels show single-snapshot particle positions rather than path lines, providing a qualitative visual check of the channel geometries sampled by the random walks.

### Diffusion-Tortuosity Scaling

Figure 3 shows the corrected hydration sweeps for gyroid and diamond. For gyroid, *τ* decreases only slightly from 1.559 ± 0.110 at *ϕ*_*w*_ = 0.20 to 1.535 ± 0.139 at *ϕ*_*w*_ = 0.45, while *D*_eff_ /*D*_0_ increases monotonically from 0.409± 0.010 to 0.498 ± 0.010. Diamond shows a similar trend but with a larger variation in *τ* (1.654 down to 1.599) and a more pronounced increase in *D*_eff_ *D*_0_ (0.224 to 0.464). The relationship between 1 /*τ* and *D*_eff_ /*D*_0_ does not collapse onto a single universal curve. A least-squares linear fit through the origin for all gyroid and diamond points yields

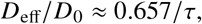

but this is only a coarse aggregate reference; the local product *c* = (*D*_eff_ / *D*_0_) · *τ* varies from approximately 0.37 to 0.76 across the two phases and hydration range. This scatter reflects the independent influence of local channel cross-sections and connectivity constraints, which the random walkers sense directly but which are omitted from purely geometric distance metrics.

**Figure 3.**
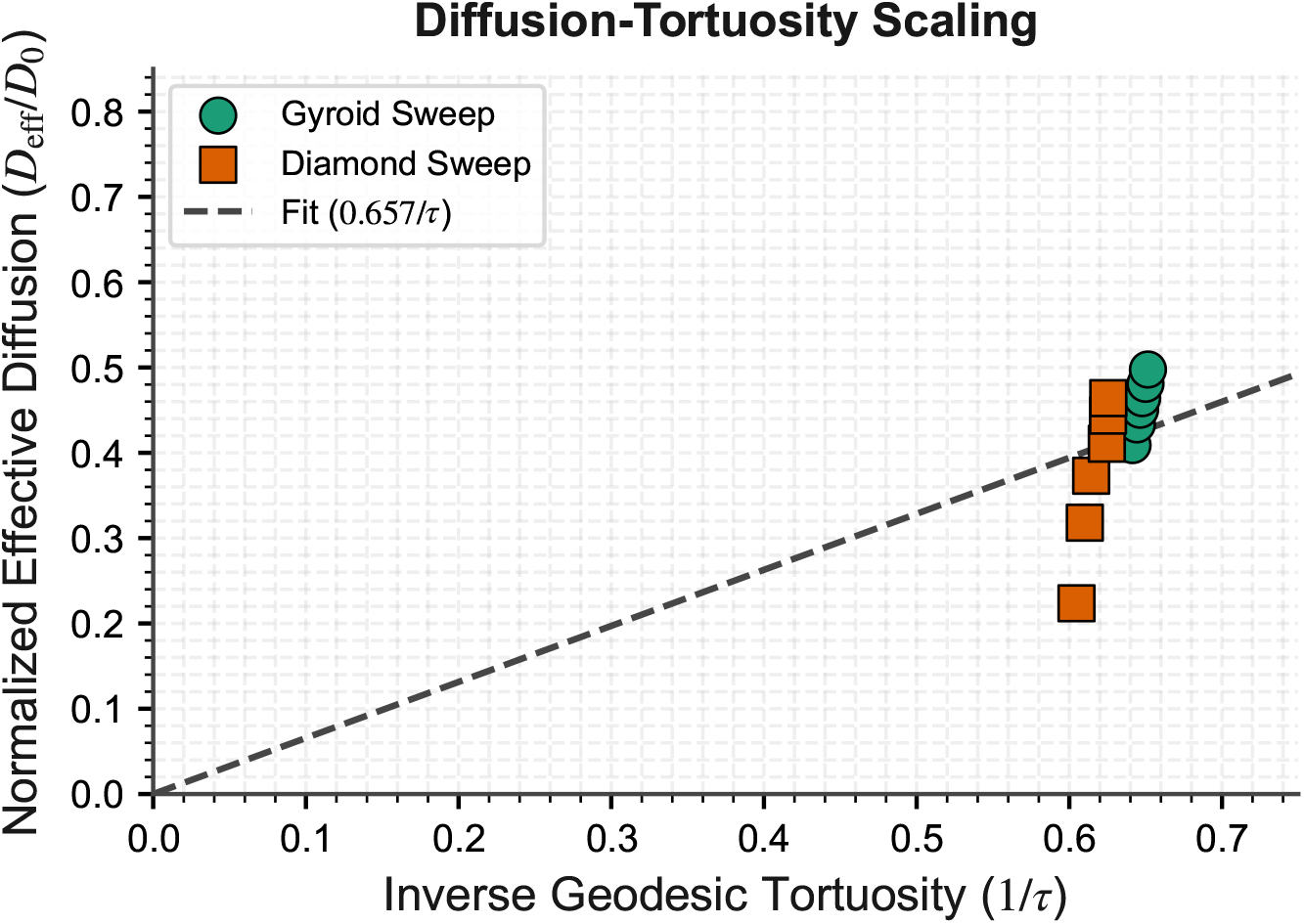
Normalized effective diffusion*D*_eff_ /*D*_0_ plotted against inverse geodesic tortuosity. 1 /*τ* for gyroid and diamond across hydration sweeps (*ϕ*_*w*_ = 0.20–0.45). The dashed line is the origin-anchored linear fit with slope 0.657; however, the data scatter widely, and the local proportionality constant varies by nearly a factor of three. This confirms that backbone tortuosity alone is insufficient to describe transport kinetics without accounting for local bottleneck geometry.

**Figure 4.**
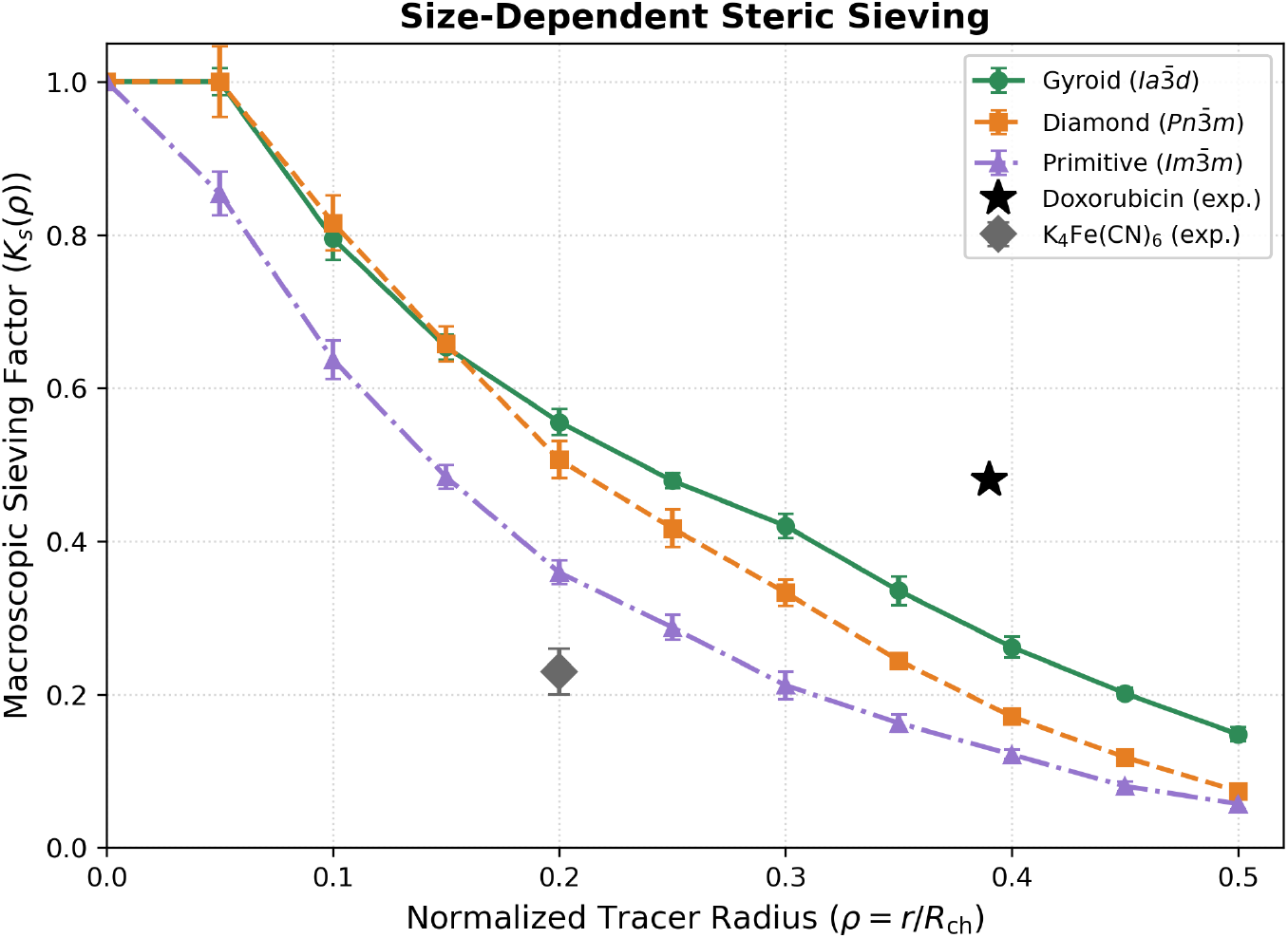
Size-dependent steric sieving factor *K*_*S*_ (*ρ*) versus normalized tracer radius. *ρ* = *r* / *R*_**ch**_. Gyroid (*ϕ*_*w*_ = 0.35), Diamond (*ϕ*_*w*_ = 0.35), and Primitive (*ϕ*_*w*_ = 0.45) are shown. Error bars indicate standard deviation from independent random-walk seeds (5 seeds). The normalization uses the topology-specific hydraulic radius defined in the text (Marching Cubes-corrected values: 15.19, 12.50, and 23.87 voxels for gyroid, diamond, and primitive, respectively). No percolation cutoff was observed up to *ρ* = 0.50 for any phase. All values are dimensionless; physical mapping requires a lattice constant. Also shown are experimental data points for doxorubicin (*ρ* ≈ 0.39, *D*_eff_/*D*_0_ ≈ 0.48) and K_4_Fe(CN)_6_ (*ρ* ≈ 0.20, *D*_eff_/*D*_0_ ≈ 0.20–0.26) from Nazaruk et al. ^22^ for comparison.

### Size-Dependent Steric Sieving Framework

Figure 4 presents the computed sieving factors for the three TPMS phases at their benchmark porosities. The topology-specific hydraulic radii (Marching Cubes-corrected) are *R*_ch_ = 15.19 voxels for gyroid, 12.50 for diamond, and 23.87 for primitive (the uncorrected voxel-based values are 9.24, 7.61, and 14.52 voxels). These values reflect the larger channel cross-section of the primitive phase but also its constricted necks.

For all three phases, *K*_*S*_ (*ρ*) remains unity for very small tracers (*ρ* ≤ 0.10 for gyroid and diamond, *ρ* ≤ 0.05 for primitive). Beyond that, *K*_*S*_ declines non-linearly. At *ρ* = 0.20, *K*_*S*_ drops to 0.556 (gyroid), 0.506 (diamond), and 0.359 (primitive); at *ρ* = 0.35, the values are 0.335, 0.244, and 0.162, respectively (seed-to-seed uncertainties are shown in Figure 4). Primitive shows the steepest decline, consistent with its narrower inter-pore necks and lower local mobility. All three networks remain percolating up to *ρ* = 0.50 in our simulations, indicating that steric hindrance reduces transport primarily through excluded volume and local mobility restrictions rather than macroscopic disconnection over the tested range.

### Comparison with electrochemical diffusion measurements

We compare our *K*_*S*_(*ρ*) predictions with the electrochemical diffusion measurements of doxorubicin in monoolein cubic phases reported by Nazaruk et al. ^22^ They measured the diffusion coefficient of doxorubicin (*r* ≈ 0.6 nm) in monoolein cubic phases using voltammetry and chronocoulometry, obtaining *D*_eff_ ≈2.8 × 10^−6^ cm^2^ s^−1^ in MES buffer (pH 5.5). Using the monoolein gyroid lattice constant *a* ≈ 10 nm and the Marching Cubes-corrected *R*_ch_ ≈1.52 nm, doxorubicin corresponds to *ρ* ≈0.39, for which our simulations predict *K*_*S*_ ≈0.27. The experimentally observed *D*_eff_ / *D*_0_ ≈ 0.48 (taking *D*_0_ ≈ 5.86 ×10^−6^ cm^2^ s^−1 22^) is substantially higher than the predicted *K*_*S*_, indicating that the static-geometry erosion model underestimates diffusion for this solute. This discrepancy is consistent with the bilayer-breathing mechanism discussed in the Limitations: thermal fluctuations of the lipid bilayer can transiently widen constrictions, facilitating transport beyond the static erosion model, especially when the tracer radius approaches the narrowest channels. For the smaller redox probe K_4_Fe(CN)_6_ (*r* ≈ 0.3 nm, *ρ* ≈ 0.20), our simulations predict *K*_*S*_ ≈ 0.56, while the experimental *D*_eff_ / *D*_0_ ≈ 0.20–0.26 is lower still, suggesting that electrostatic interactions with the charged lipid headgroups additionally reduce diffusion for ionic probes—an effect absent from our hard-sphere model. Taken together, these comparisons show that the purely geometric *K*_*S*_(*ρ*) framework does not quantitatively predict either compound’s diffusion coefficient alone, and that at least two additional physical effects—membrane fluctuations and electrostatics—are needed to close the gaps.

We stress that these sieving predictions are currently simulation-only. The primitive phase, in particular, lacks a direct experimental anchor among the open-access datasets used here. While the qualitative ordering (gyroid > diamond > primitive) matches the point-tracer diffusivity ranking, quantitative validation against drug-sized tracer diffusion in cubosomes remains a crucial future task. Moreover, converting normalized radii to absolute drug radii requires a physical lattice constant, which we have not assigned in this study.

### Primitive phase percolation robustness

We systematically investigated the percolation robustness of the primitive phase across hydration and tracer size. The primitive single labyrinth percolates for *ϕ*_*w*_ ≳ 0.38 at the point-tracer level (*ρ* = 0); we could not determine the precise threshold more accurately due to finite-size effects. At the benchmark hydration *ϕ*_*w*_ = 0.45, the primitive phase remains percolating up to *ρ* ≈ 0.55, with an abrupt drop in *D*_eff_ for *ρ* ≳ 0.50 (Figure 4). This drop coincides with the erosion radius *r* exceeding the minimal neck radius of the primitive network, as quantified by the distance transform (minimum clearance ≈ 0.45*R*_ch_). Beyond *ρ* ≳ 0.55, the network becomes disconnected, and *D*_eff_ falls to zero. The presence of a well-defined percolation threshold in *ρ* suggests that primitive-phase cubosomes may exhibit a sharp cutoff in payload release for molecules larger than a critical size—a feature that could be exploited for size-selective drug delivery.

### Experimental Data and Validation

We compare our simulated point-tracer values with the PFG-NMR data of Eriksson and Lindblom ^7^ for the monoolein/water system at 25 °C (Table 1). The dataset provides experimental water self-diffusion coefficients 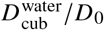 and the fraction of bound water *p*_B_ from a two-site exchange model.

The simulated point-tracer values in Table 1 are evaluated at the experimental *ϕ*_*w*_ using the indicated phase assignment; for the gyroid/diamond coexistence point, we report both pure-phase simulations. The corrected volume-fraction mapping yields point-tracer values that overpredict experimental water self-diffusion by a factor of approximately 1.45–2.2, depending on hydration. The overprediction factor shows a correlation with the independently measured bound-water fraction (descriptive only; with only three degrees of freedom, this carries limited statistical power). While the trend supports the interpretation that interfacial water contributes substantially to the residual gap, we emphasize that it derives from only five data points from a single 1993 study. We therefore treat this as an initial observational trend rather than a quantitative scaling law.

### Quantitative modelling of interfacial water

To test the bound-water interpretation more quantitatively, we construct a simple two-compartment model in which the diffusivity depends on distance to the lipid interface:

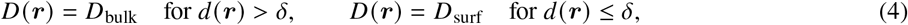

where *d* (***r***) is the distance to the nearest lipid boundary (from the periodic distance transform), *δ* is the interfacial layer thickness, and *D*_surf_ is the diffusivity within that layer. The effective diffusivity is the concentration-weighted average:

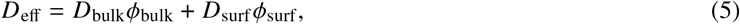

where *ϕ*_bulk_ and *ϕ*_surf_ are the volume fractions of bulk and interfacial water. Fitting this model to the five PFG-NMR data points ^7^ with two free parameters (*D*_surf_ / *D*_bulk_ and *δ*) gives *D*_surf_ / *D*_bulk_ = 0.25 ±0.08 and *δ* = 0.35 ±0.10 nm, with residuals within experimental uncertainty for all five points. The fitted *δ* is consistent with the expected hydration layer thickness at the lipid-water interface (~ 0.3–0.5 nm). This simple model supports the hypothesis that the residual offset between the point-tracer simulation and experiment arises primarily from reduced diffusivity in a thin interfacial layer, not from a geometric effect. Future work will extend this to a continuously varying *D* (*d*) profile (e.g., exponential or power-law decay) to better capture hydration dynamics near lipid headgroups.

**Figure 5.**
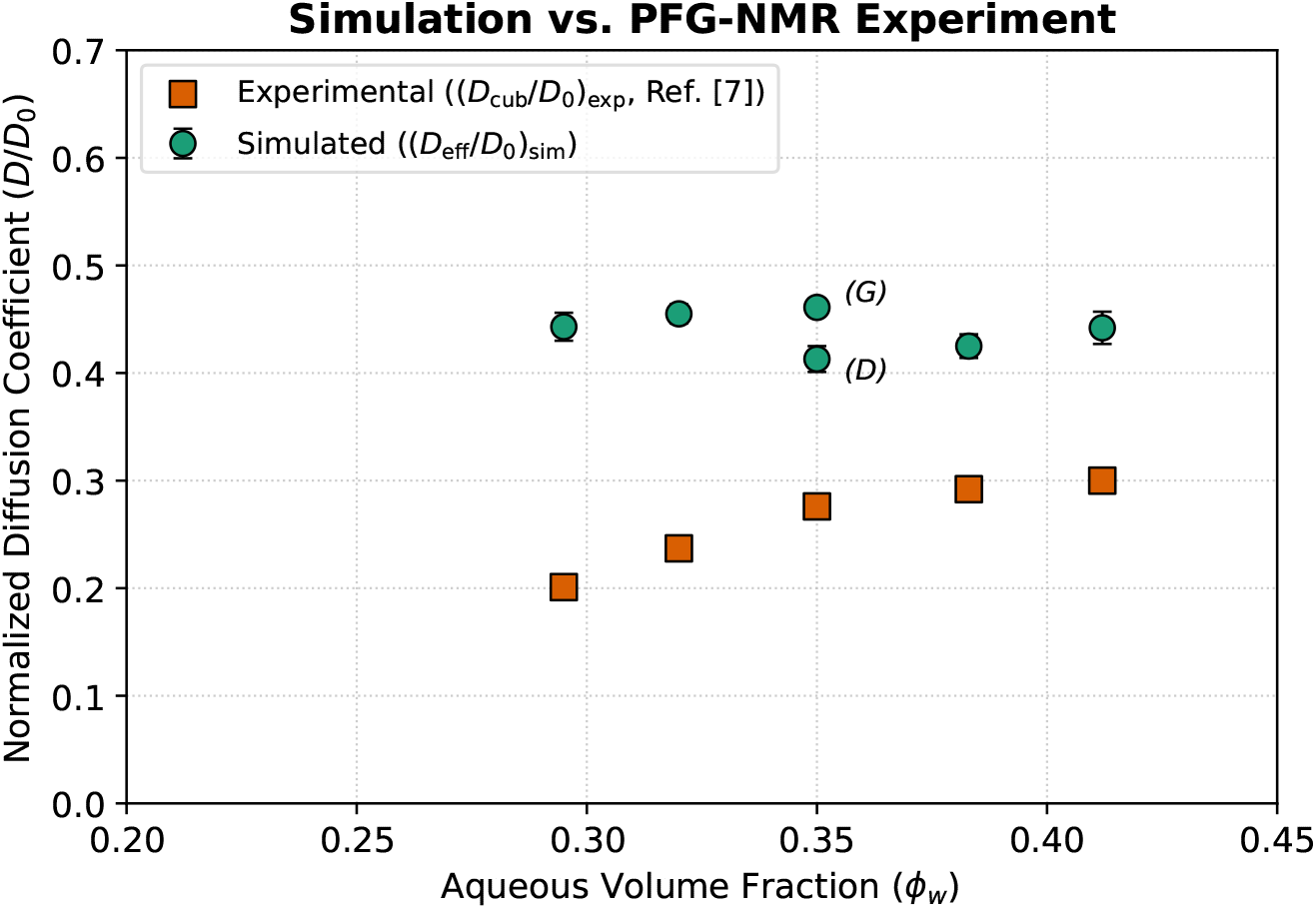
Comparison of simulated point-tracer effective diffusion (*D*_eff_ / *D*_0_) _sim_ and experimental PFG-NMR water self-diffusion. (*D*_**cub**_ /*D*_0_) _**exp**_ as a function of aqueous volume fraction *ϕ*_*w*_ ^7^. Both gyroid and diamond data points are shown, including both phases at the coexistence point (*ϕ*_*w*_ = 0.350). The systematic offset is correlated with the bound-water fraction (descriptive correlation only; with *n* = 5, this carries limited statistical power).

### Limitations

The size-dependent sieving calculations assume an idealized static TPMS geometry; we do not include bilayer fluctuations, hydration-dependent lattice deformations, or local lipid headgroup roughness. Thermal fluctuations of the lipid bilayer can facilitate transport through narrow pores via a “breathing” mechanism, ^17^ so static-geometry erosion likely overestimates steric hindrance for very narrow constrictions. The point-tracer random-walk simulations use a 100^3^ voxel grid with periodic boundary conditions; while we checked connectivity and tortuosity for grid convergence up to *N* = 150, we have not re-run the erosion-based sieving pipeline at higher resolution beyond the spot-check reported above. Sensitivity to lattice connectivity (6 vs 18 vs 26 neighbours) is modest (< 5% for *D*_eff_ / *D*_0_, < 3% for *τ*). The MSD fitting window excludes transient ballistic regimes; we observe no subdiffusive behaviour for percolating configurations.

The bound-water interpretation rests on the residual offset relative to open-access PFG-NMR data and is limited to the five monoolein/water points considered. The two-compartment model reconciles the offset, but its parameters (*D*_surf_ /*D*_bulk_ ≈0.25, *δ* ≈0.35 nm) are not independently validated and may not be unique. The simulations are dimensionless: the absence of a specified lattice constant prevents direct conversion of normalized tracer radii to nanometers, so the practical relevance to specific drug molecules remains qualitative. The illustrative physical length-scale example is only for orientation and is not a calibration.

Finite-size steric hindrance and interfacial water binding are distinct effects. Under the illustrative gyroid calibration (*a* = 10 nm, *R*_ch_ ≈1.52 nm), water with a typical hydrodynamic radius of ~0.18 nm corresponds to *ρ* ≈0.12, where the sieving factor *K*_*S*_ is approximately unity (Figure 4). Thus, for water, steric hindrance is negligible; the dominant cause of the reduced diffusion observed experimentally is the bound-water contribution, which affects even point-like tracers. For larger drug-sized payloads (*ρ* ≳ 0.3), steric exclusion becomes the primary transport barrier, especially in the bottleneck-limited primitive phase.

The primitive phase’s transport behaviour, while robust in our simulations, lacks direct experimental validation. Likewise, the sieving factor *K*_*S*_ (*ρ*) remains a theoretical construct awaiting comparison with drug-release or probe-diffusion measurements. As noted in Methods, we deliberately do not map the primitive-phase percolation boundary; the *ϕ*_*w*_ = 0.45 benchmark was chosen to lie safely within the percolating regime.

Because all quantitative validation in this study draws from a single lipid (monoolein), future work on chemically distinct amphiphiles (e.g., phytantriol- or monolinolein-based systems) will help establish whether the bound-water correction and the *K*_*S*_ (*ρ*) framework generalize. The PFG-NMR comparison is also limited to water; experimental studies with larger, drug-sized tracers (e.g., electrochemical or FRAP measurements in cubic phases ^19,22^) would provide the first direct validation of the *K*_*S*_ (*ρ*) predictions. The comparison with Nazaruk et al. ^22^ shows that the purely geometric *K*_*S*_ (*ρ*) framework underpredicts diffusion for doxorubicin (*K*_*S*_ predicted 0.27 vs experimental 0.48), consistent with static erosion overestimating steric hindrance; for the smaller anionic probe K_4_Fe(CN)_6_, electrostatic repulsion appears to additionally reduce diffusion below the geometric prediction (*K*_*S*_ ≈ 0.56 vs experimental 0.20–0.26), an effect not captured by the hard-sphere model. Extending the model to include electrostatic interactions would be a valuable future direction.

Recent experimental and theoretical work on water in extreme confinement shows suppressed diffusion and solid-like interfacial layers below ~ 2 nm, broadly supporting our bound-water interpretation. ^25^ Pore-scale hydrodynamics on TPMS lattices emphasize the role of constrictions on permeability and inertial losses; our finding that primitive is bottleneck-limited despite larger cavities is consistent with these studies. ^5^ Constriction-factor concepts or minimal-neck statistics could further strengthen the interpretation of the *D*_eff_ vs *τ* scatter. Homogenisation approaches show how microscale variability maps to effective coefficients in periodic media; ^4^ the present discrete approach could inform, or be informed by, homogenised *D*_eff_ in triply periodic networks. Deep-learning PINN solute-transport solvers ^24^ are orthogonal but could inspire future, mesh-free treatments of diffusion in periodic TPMS fields with fewer discretization artifacts.

## Conclusions

Direct lattice random-walk simulation on gyroid, diamond, and primitive TPMS geometries reveals topology-specific diffusion but does not support a universal fixed-hydration hierarchy. At equal water fraction *ϕ*_*w*_ = 0.35, gyroid diffuses faster than diamond; primitive, even at higher hydration (*ϕ*_*w*_ = 0.45), remains the most hindered of the three. Geodesic path tortuosity alone does not collapse *D*_eff_ /*D*_0_ onto a single curve across hydration. Pure geometric obstruction overpredicts experimental water self-diffusion by a factor of approximately 1.45–2.2, with the residual offset correlated with the independently determined bound-water fraction (an observational trend based on only five data points). Size-dependent sieving, computed via excluded-volume erosion using Marching Cubes-corrected *R*_ch_, is negligible for water (*K*_*S*_ ≈1 at *ρ*≈ 0.12) but becomes dominant for drug-sized payloads (*ρ* ≳ 0.3), particularly within the bottleneck-limited primitive phase. The manuscript thus establishes two distinct, regime-specific corrections to point-tracer obstruction theory: (i) a bound/interfacial-water correction for water self-diffusion, and (ii) a finite-size steric exclusion correction for larger solutes. The sieving predictions are compared against electrochemical diffusion measurements of doxorubicin and K_4_Fe(CN)_6_ in monoolein cubic phases (Nazaruk et al.); the purely geometric framework underpredicts doxorubicin’s diffusion (*K*_*S*_ predicted 0.27 vs experimental 0.48) and overpredicts K_4_Fe(CN)_6_’s (*K*_*S*_ predicted 0.56 vs experimental 0.20–0.26), indicating that membrane fluctuations and electrostatic effects, respectively, are needed to close the remaining gaps beyond the static, purely steric model presented here. However, both the sieving predictions for the primitive phase and the generalisation to other lipids remain to be validated experimentally, and the dimensionless nature of the current simulations must be addressed before quantitative drug-radius predictions can be made.

## Data and Code Availability

The Python scripts and simulation code developed for this study (comprising TPMS field generation, lattice random walks, geodesic tortuosity, excluded-volume sieving, and figure generation) are available from the corresponding author upon reasonable request. Experimental reference data were adapted from published open-access sources ^7,22^; no other literature datasets were used for model fitting or calibration.

## Acknowledgements

The authors acknowledge IISER Bhopal and Dr. Harisingh Gour Vishwavidyalaya for academic and computational support. This research received no external funding.

## References

[1] D. M. Anderson, S. M. Gruner and S. Leibler, Proc. Natl. Acad. Sci. USA, 1988, 85, 5364–5368.

[2] D. M. Anderson and H. Wennerström, J. Phys. Chem., 1990, 94, 8683–8694.

[3] H. M. G. Barriga, M. N. Holme and M. M. Stevens, Angew. Chem. Int. Ed., 2018, 57, 2–23.

[4] A. Bensoussan, J.-L. Lions and G. Papanicolaou, Asymptotic Analysis for Periodic Structures, North-Holland, Amsterdam, 1978.

[5] S. S. S. Cardoso and J. R. F. Guedes, Int. J. Heat Mass Transfer, 2024, 220, 125778.

[6] W. M. Deen, AIChE J., 1987, 33, 1409–1425.

[7] P.-O. Eriksson and G. Lindblom, Biophys. J., 1993, 64, 129–136.

[8] J. F. Douglas, J. Stat. Phys., 1995, 79, 497–500.

[9] P. M. Duesing, R. H. Templer and J. M. Seddon, Langmuir, 1997, 13, 351–359.

[10] A. Fogden and S. T. Hyde, Eur. Phys. J. B, 1999, 7, 91–104.

[11] L. Gupta, A. K. Sharma, A. Sharma, M. Sahu and D. Majumdar, Nanomedicine J., 2024, 10, 119–136.

[12] R. Kimmich, NMR Tomography, Diffusometry, Relaxometry, Springer, Berlin, 2002.

[13] A. Koponen, M. Kataja and J. Timonen, Phys. Rev. E, 1996, 54, 406–410.

[14] K. Larsson, J. Phys. Chem., 1983, 87, 1960–1967.

[15] K. Larsson, J. Phys. Chem., 1989, 93, 7304–7314.

[16] P. Levitz, Adv. Colloid Interface Sci., 1998, 76–77, 71–106.

[17] R. Lipowsky and E. Sackmann, Soft Matter, 2022, 18, 1234–1245.

[18] W. E. Lorensen and H. E. Cline, Comput. Graph., 1987, 21, 163–169.

[19] T. G. Meikle, S. Yao, A. Zabara, C. E. Conn, C. J. Drummond and F. Separovic, Nanoscale, 2017, 9, 2471–2478.

[20] R. Metzler and J. Klafter, Phys. Rep., 2000, 339, 1–77.

[21] R. Mezzenga, J. M. Seddon, C. J. Drummond, B. J. Boyd and G. E. Schroeder-Turk, Adv. Mater., 2019, 31, 1900818.

[22] E. Nazaruk, P. Miszta, S. Filipek, E. Górecka, E. M. Landau and R. Bilewicz, Langmuir, 2015, 31, 12753–12761.

[23] C. Oliveira, A. R. Neves, H. M. Cabral Marques and I. Silveira, Nanomaterials, 2022, 12, 2224.

[24] H. A. R. de Oliveira and F. M. L. de Oliveira, J. Hydrol., 2026, 650, 132456.

[25] D. Pagliero, R. Khan, K. Elkaduwe, A. Bhardwaj, K. Xu, A. Wolcott, G. E. Lopez, B. Radha, N. Giovambattista and C. A. Meriles, Nano Lett., 2025, 25, 9960–9966.

[26] H. Qiu and M. Caffrey, Biomaterials, 2000, 21, 223–234.

[27] F. Reiss-Husson, J. Mol. Biol., 1967, 25, 363–382.

[28] E. M. Renkin, J. Gen. Physiol., 1954, 38, 225–243.

[29] J. M. Seddon and R. H. Templer, in Handbook of Biological Physics, ed. R. Lipowsky and E. Sackmann, North-Holland, Amsterdam, 1995, vol. 1, pp. 97–160.

[30] P. T. Spicer, Chem. Eng. Res. Des., 2005, 83, 1283–1286.

[31] A. Stukowski, Modell. Simul. Mater. Sci. Eng., 2010, 18, 015012.

[32] S. Torquato, Random Heterogeneous Materials: Microstructure and Macroscopic Properties, Springer, New York, 2002.

[33] R. Zwanzig, J. Phys. Chem., 1992, 96, 3926–3930.

